# Lipid analysis of CO_2_-rich subsurface aquifers suggests an autotrophy-based deep biosphere with lysolipids enriched in CPR bacteria

**DOI:** 10.1101/465690

**Authors:** Alexander J. Probst, Felix J. Elling, Cindy J. Castelle, Qingzeng Zhu, Marcus Elvert, Giovanni Birarda, Hoi-Ying Holman, Katherine R. Lane, Bethany Ladd, M. Cathryn Ryan, Tanja Woyke, Kai-Uwe Hinrichs, Jillian F. Banfield

## Abstract

Sediment-hosted CO_2_-rich aquifers deep below the Colorado Plateau (USA) contain a remarkable diversity of uncultivated microorganisms, including Candidate Phyla Radiation (CPR) bacteria that are putative symbionts unable to synthesize membrane lipids. The origin of organic carbon in these ecosystems is unknown and the source of CPR membrane lipids remains elusive. We collected cells from deep groundwater brought to the surface by eruptions of Crystal Geyser, sequenced the community, and analyzed the whole community lipidome over time. Characteristic stable carbon isotopic compositions of microbial lipids suggest that bacterial and archaeal CO_2_ fixation ongoing in the deep subsurface provides organic carbon for the complex communities that reside there. Coupled lipidomic-metagenomic analysis indicates that CPR bacteria lack complete lipid biosynthesis pathways but still possess regular lipid membranes. These lipids may therefore originate from other community members, which also adapt to high in situ pressure by increasing fatty acid unsaturation. An unusually high abundance of lysolipids attributed to CPR bacteria may represent an adaptation to membrane curvature stress induced by their small cell sizes. Our findings provide new insights into the carbon cycle in the deep subsurface and suggest the redistribution of lipids into putative symbionts within this community.

## INTRODUCTION

The most prominent characteristic of the deep continental subsurface is the absence of sunlight. However, the diversity of subsurface ecosystems is manifold. Physicochemical characteristics as well as the availability of electron donors and acceptors shape different microbial communities within these ecosystems (e.g., [1, 2]). Serpentinizing systems can harbor large amounts of abiotically-produced methane that can serve as the primary electron donor and as a carbon substrate entering biosynthesis via methanotrophy [3–5]. In some environments, including petroleum deposits, the availability of fossil organic matter, burial depth, and temperature, may exert strong control on community structure [6]. Other subsurface environments have low availability of buried organic matter. In such environments, genomic analyses suggest that *in situ* CO_2_ fixation may support microbial communities [7–9]. Most subsurface environments may be sustained by fixed carbon from multiple sources, and the relative importance of *in situ* CO_2_ fixation has been difficult to ascertain [10].

The candidate phyla radiation (CPR) of bacteria is a monophyletic group [11] which includes enigmatic small-celled microbes [12] that appear to be abundant predominantly in the subsurface [13]. Co-cultures of CPR bacteria indicate that some are symbionts of other bacteria and heavily depend on their hosts for basic resources [14]. To date, none of the reconstructed CPR genomes encode for a complete fatty acid-based lipid biosynthesis pathway [13]. Other bacterial and archaeal symbionts from different branches of the tree of life also do not encode for their own lipid biosynthesis pathway [15–17] and at least one hyperthermophilic episymbiont (*Nanoarchaeum equitans*) has been suggested to acquire its lipids from the host archaeon [18]. However, the origin and types of lipids used by CPR bacteria remain elusive.

Analysis of the stable carbon isotopic ratios of lipid molecules has enabled researchers to track carbon flow through communities in cases where the communities were very simple. For instance, it was shown that archaea growing in syntrophy with sulfate-reducing bacteria mediate the anaerobic oxidation of methane [19, 20]. This analysis was possible because the consortia were based on simple bacterial and archaeal assemblages that produce diagnostic lipid types. In another study, the stable carbon isotope ratios of methane and lipids were used to track the flow of carbon from methane into the two species thought to be present based on rRNA sequence profiling [21]. The power of this approach is limited when microbial communities are complex and contain numerous organisms that produce unknown lipid molecules. In fact, lack of information about the types of lipids produced by uncultivated organisms remains a major gap in microbial ecology.

A recent large-scale environmental genomics survey of subsurface microbial ecosystems within the Colorado Plateau, USA, provided evidence for a depth-based distribution of organisms affiliated with more than one hundred different phylum-level lineages [10]. Samples were acquired from groundwater that erupted through the cold, CO_2_-driven Crystal Geyser. During the eruption cycle groundwater was sourced from different depths, enabling the assignment of organisms to their respective aquifer regions. Genomic resolution of the tracked organisms linked three different carbon fixation pathways to groundwater from different depths. However, a major question remains regarding the extent to which autotrophic organisms provide organic carbon to these complex microbial communities. Further, the types and sources of lipids used to construct the cell envelope of CPR bacteria remain elusive. We postulated that clues regarding the types of lipids produced by uncultivated bacteria and archaea could be addressed by correlation-based analyses so long as sufficient numbers of samples were defined in terms of the abundances of the microorganisms present and overall lipid compositions of the same samples were available. Here, we use coupled metagenomic-lipidomic datasets to test this approach and to resolve the importance of autotrophy as the source of organic carbon in the studied environment.

## MATERIAL AND METHODS

### Sampling scheme

Samples for lipid analyses were retrieved by collecting cells in groundwater from the Crystal Geyser ecosystem onto a 0.1-µm teflon filter (Gravertech 10” MEMTREX-HFE). Cells were immediately frozen on dry ice. One post 0.2-µm fraction was also collected to enrich for organisms of the CPR and DPANN radiations. The samples span an entire cycle of the geyser, which lasted for approximately 5 days [10]. Collection for each metagenomic sample proceeded for around four hours. Collection of lipid samples proceeded simultaneously, but the collection time was around 8 hours so there are half as many lipid samples as metagenome samples. The sampling scheme details are presented in Fig. S1.

### Sampling and isotopic analysis of dissolved inorganic carbon

Twenty-four groundwater samples were collected from about 8.5 m below ground surface in the geyser borehole using a peristaltic pump and copper pipe. Samples were collected in 12 mL glass vials. The vials were flushed with fresh geyser water and were filled underwater in a bucket that was overflowing with groundwater to avoid atmospheric contact; this was confirmed by gas chromatography analyses that had no detectable N_2_, O_2_, or Ar concentrations, and 100% CO_2_ composition (unpublished data). The stable carbon isotopic composition of the dissolved organic carbon was analyzed by Continuous Flow Isotope Ratio Mass Spectrometry (CF-IRMS) using a Thermo Finnigan GasBench coupled to a DeltaV^Plus^. Water pressure, temperature, and electrical conductivity were measured in situ at the same depth using a Solinst LTC Levelogger Edge.

### Lipidomics

Methods for lipid extraction and analysis are described in detail in the supplementary information. In brief, lipids were extracted using a modified Bligh and Dyer method [22] after addition of an internal standard. Archaeal and bacterial intact polar lipids (IPLs; for structures see Fig. S2) were quantified using a Dionex Ultimate 3000 ultra-high performance liquid chromatography (UPLC) system connected to a Bruker maXis Ultra-High Resolution quadrupole time-of-flight mass spectrometer equipped with an electrospray ion source operating in positive mode (Bruker Daltonik, Bremen, Germany). Lipids were separated using normal phase UPLC on an Acquity UPLC BEH Amide column (1.7 µm, 2.1 × 150 mm; Waters Corporation, Eschborn, Germany) maintained at 40 °C as described in ref. [23]. For isotopic analysis, IPLs were separated from free core lipids using semi-preparative high performance liquid chromatography. For mass spectrometric analysis of previously uncharacterized IPLs see Figs. S3 and S4. Ether cleavage and saponification were performed on the IPL fractions to release isoprenoid hydrocarbons and fatty acids, respectively. The stable carbon isotopic compositions of these compounds were analyzed using gas chromatography–IRMS. Fourier-transform infrared (FTIR) spectromicroscopy was performed to detect lipids in intact cells. The FTIR system consisted of a Hyperion 3000 Infrared-Visible microscope coupled to a Vertex70V interferometer (Bruker Optics – Billerica MA). For FTIR analysis, cells were deposited on a double-side-polished silicon slide and dried with a gentle nitrogen gas stream in a biological safety cabinet. Lipid identification was achieved by comparing spectra from samples and dry films of lipid standards.

### Metagenomics

Methods for DNA extraction, and metagenomic sequencing are described in ref. [10]. In brief, DNA was extracted from filters using the MoBio PowerMax Soil DNA isolation kit, and library preparation and sequencing were performed at the Joint Genome Institute (details on extracted DNA, type of library and sequencing are provided in ref. [10]). Quality filtered reads (https://github.com/najoshi/sickle, https://sourceforge.net/projects/bbmap) were assembled using IDBA_UD [24], genes were predicted using prodigal (meta-mode; [25]). Coverage of scaffolds was calculated using bowtie2 (–sensitive) [26]. Taxonomy of scaffolds was determined by searching proteins against an in-house database.

### Tracking taxa across time using ribosomal protein S3

In order to get a near complete picture of specific taxa present in the samples, we extracted ribosomal protein S3 (rpS3) sequences from all assembled scaffolds >1 kb using separately designed HMMs for archaea, bacteria and eukaryotes (https://github.com/AJProbst/rpS3_trckr). The extracted amino acid sequences were clustered at 99% identity (collapsing most of the strains of the same species [27]) and the longest scaffold bearing a representative rpS3 sequence was obtained for each cluster. Using read mapping (bowtie2, [26]) and allowing a maximum of three mismatches per read (according to the 99% identity of the de-replicated rpS3 sequences), the relative abundance of each selected rpS3 scaffold was calculated across all samples. The breadth (i.e. how much of the sequence of a scaffold is covered) of the scaffolds was calculated in each sample. To call a rpS3 sequence present in a sample, it had to be either assembled or have a breadth of at least 95% of the entire scaffold in a sample. Since we worked with scaffolds, we did not consider ambiguous bases for calculating the breadth. The rpS3 sequences were taxonomically annotated against a combined database from previous publications [10, 28, 29], which was de-replicated at 99% rpS3 identity. Taxonomic assignments were performed with similarity cutoffs as described earlier: ≥99% for species, ≥95% for genus, ≥90% for family level. Lower percentages were assigned to phylum or domain level (<50%).

### Statistical analysis to correlate taxa abundance with IPLs

Relative abundance measures of rpS3 genes were correlated (Pearson correlation) with relative abundance measure of IPLs if the rpS3 gene / the IPL species was present in at least 7 out of 14 samples. Resulting p-values underwent false discovery correction using the Bonferroni procedure and these q-values were then weighted by division of the q-value with the percent relative abundance of the rpS3 gene. Each lipid was allowed to be assigned to only one organism (with the best score). This assignment of lipids to rpS3 genes considers that highly abundant organisms are more likely to be detected in lipid analyses than low abundant organisms. IPL signatures were co-correlated (Bonferroni-corrected p-value <0.005) and lipid species that correlated with other lipids were identified for further analyses. These co-correlated lipid species as well as the correlation of rpS3 genes and lipid species were used to construct a network. Primary lipids were assigned based on direct correlation of lipids with organisms, secondary lipids were assigned based on a correlation with primary lipids. Lipids were classified as unspecific if the secondary lipid correlated with two primary lipids of different organisms.

### Binning of genomes

rpS3 genes that were not found in existing genomes [10, 30] were identified based on a similarity (<98% [31]) and searched for in the respective metagenomes. Genomes containing these rpS3 sequences were binned using a consensus of guanine-cytosine content, coverage and taxonomy information in the ggKbase platform [32]. Genomes were subsequently curated with ra2 [33] for scaffolding errors.

### Genomic analysis of lipid biosynthesis pathways in CPR genomes

Protein sequences were annotated from USEARCH (–ublast) searches against UniProt, UniRef100 [34], and KEGG databases [35] and uploaded to ggKbase (https://ggkbase.berkeley.edu). Based on existing annotations target proteins involved in bacterial fatty acids, isoprenoids and lipids biosynthesis were identified in CPR genomes and can be accessed using the following link: https://ggkbase.berkeley.edu/genome_summaries/1491-Bacterial_membrane_lipids_AJP.

## RESULTS AND DISCUSSION

### Microbial community profile based on marker genes

We *de novo* assembled 27 metagenome samples, the reads from which were previously used in a study that involved mapping to 505 genomes reconstructed from prior datasets to link organisms to groundwater of different depths [10]. In the current study, we extracted assembled sequences of ribosomal protein S3 (rpS3) and used read mapping to scaffolds carrying this gene to follow organisms over the 5-day eruption cycle. This approach allowed us to track 914 putatively distinct microbial species (Fig. 1), greatly exceeding the 505 previously reconstructed genomes [10].

**Fig. 1:**
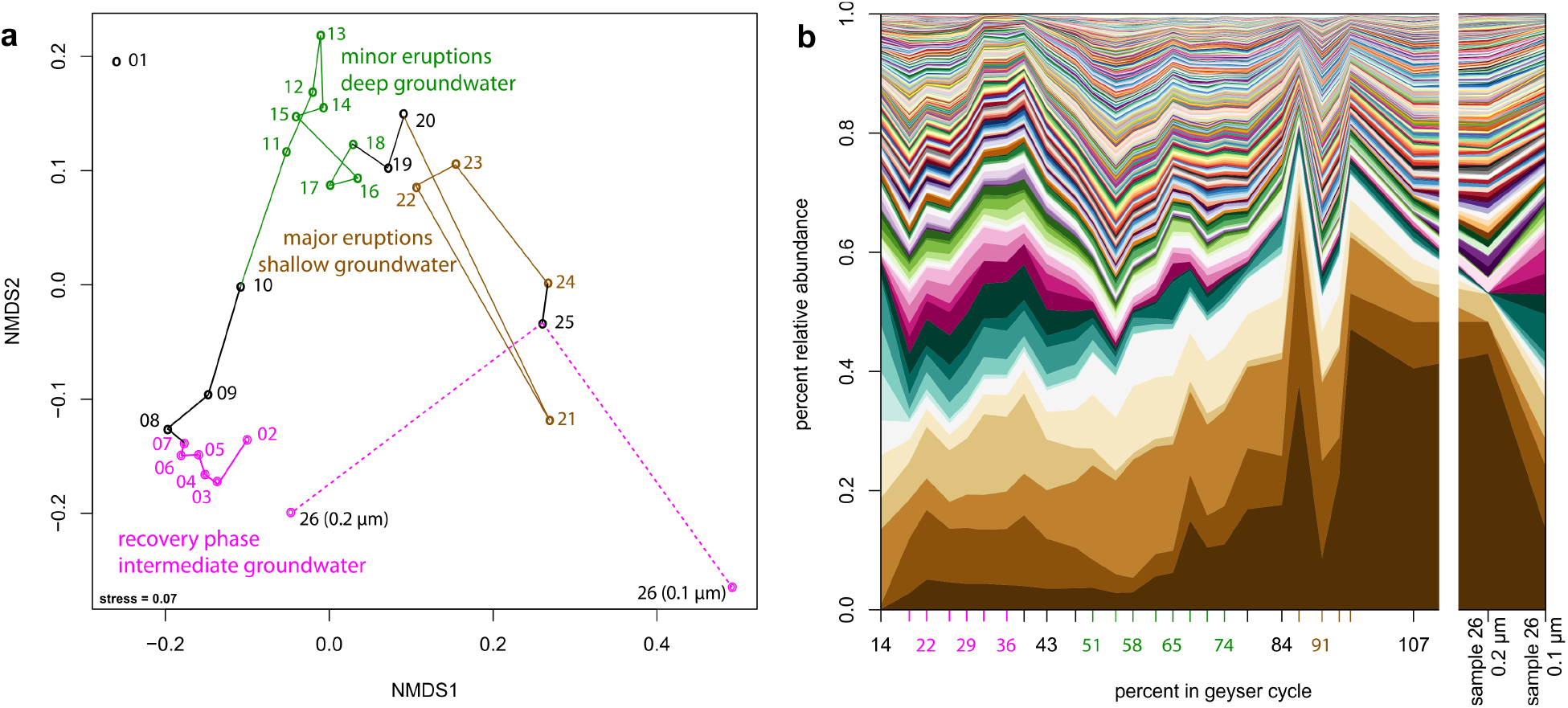
Community structure of 27 metagenomic samples from Crystal Geyser based on percent relative abundance of scaffolds carrying rpS3 sequences (clustered at 99% amino acid similarity). **A**. Non-metric multidimensional scaling based on the Bray-Curtis index. The connections show the trajectory of the different samples taken throughout the eruption cycle. Sample 01 was not included as it was an amplified library due to low biomass (see Material and Methods for further details). **B**. Relative abundance of different rpS3-carrying scaffolds across the different samples (based on stringent read-mapping). Each sample is one line on the x-axis One color represents one microbial rpS3 gene that was tracked over time.

We detected a large community shift associated with transition to different eruption phases. According to previously published geochemical data [10], the first phase, referred to as the recovery phase, sources groundwater from an aquifer of intermediate depth, likely a Navajo Sandstone-hosted aquifer. During the second minor eruption phase water from a deeper aquifer is sourced (likely Wingate Sandstone-hosted) and during the third major eruption phase an increased fraction of shallow groundwater is sourced (Fig. 1). Grouping of samples into different clusters in an ordination analysis based on community composition (Fig. 1A) revealed stepwise changes throughout the eruption cycle. The final sample, which was taken after the end of the major eruption phase and as the geyser transitions into the next recovery phase was size-fractionated, with cells collected sequentially on a 0.2 µm filter and followed by a 0.1 µm filter (sample 26, Fig. 1). The community composition on the 0.2 µm filter plots near samples from the beginning of the first cycle in the ordination analysis, indicative of a restoration of the initial microbial community (Fig. 1A).

### *In situ* carbon fixation sustains microbial communities irrespective of aquifer depth

Previous community-wide genomic analyses suggested that carbon fixation might sustain the relatively complex aquifer microbial communities, but direct evidence was lacking. We measured the stable carbon isotope composition (that is, δ^13^C values) of IPL-derived bacterial fatty acids (FA) and archaeal phytane. The values for 14 samples were plotted as a function of sampling time and compared to the δ^13^C values of DIC and CO_2_ in the ecosystem (Fig. 2A, B). The δ^13^C values for DIC sampled from the geyser discharge over its five-day cycle ranged from 3.6 to 8.0‰ (average = 5.0‰, std. dev. = 1.4‰) and showed no systematic variation with relative depth of source water (Fig. S5). The δ^13^C values for phytane range between -47.0 and -32.8‰ and for bacterial lipids (expressed as weighted average of all FAs) from -32.7 to -22.1‰. We found very little genomic evidence for utilization of methane [30] by these communities and methane was not detected in the geyser gas emissions [10]. Thus, we do not attribute the ^13^C-depletion of phytane to methane metabolism by methanogens/methanotrophs. The Wingate and Navajo aeolian sandstone aquifers have little associated sedimentary organic carbon [36, 37]. Thus, the ^13^C-depletion in lipids likely does not originate from heterotrophic incorporation of sediment-associated organic carbon.

**Fig. 2:**
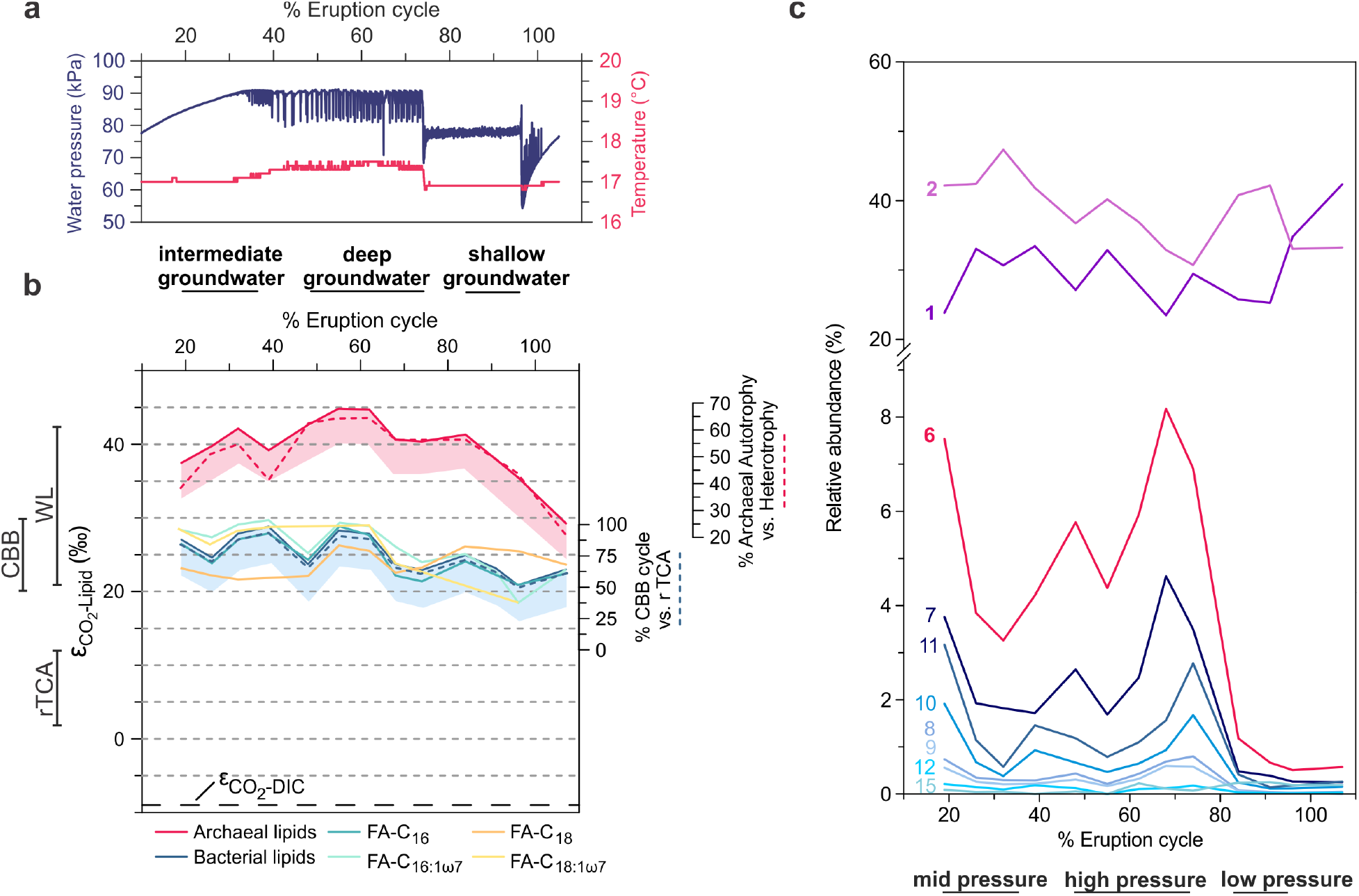
Carbon isotopic ratios and relative abundance of unsaturated intact polar lipids relative to the cycle of the geyser. **a)** Water pressure and temperature over the geyser cycle showing sourcing of fluids from the conduit (mixed), the deep aquifer, and the shallow aquifer from ref. [10]. **b**) Stable carbon isotope fractionation of archaeal lipids (phytane, released from archaeol), individual bacterial fatty acids (FA, released from diacylglycerols), bacterial lipids (weighted average of FA), and dissolved inorganic carbon (DIC) relative to CO_2_ (εCO_2_-Lipid) over the geyser cycle. Lines to the left of the panel show expected ranges of εCO_2_-Lipid (assuming εCO_2-_Lipid=εCO2-Biomass, potential depletion of bacterial and archaeal lipids relative to biomass by ~5‰ is indicated by shaded areas) for the Calvin-Bassham-Benson (CBB; [39, 42, 43]), the reductive tricarboxylic acid cycle (rTCA [39, 44, 45]), and the Wood–Ljungdahl pathway (WL, reductive acetyl-coenzyme A pathway; [39, 45, 46]). The blue dashed line indicates relative contribution of carbon fixation through the CBB cycle versus the rTCA cycle for bacterial lipids (assuming maximum fractionation due to high in situ [CO_2_] and [DIC]). The red dashed line indicates the relative contribution of autotrophy versus heterotrophy (uptake of bacterial CBB/rTCA-fixed carbon) to archaeal lipid biomass, calculated from mass balance of δ^13^C values of bacterial and archaeal lipids (assuming maximum fractionation for archaeal autotrophy due to high in situ [CO_2_] and [DIC]). **c)** Relative abundance of unsaturated diacylglycerol membrane lipids (the number indicates the sum of double bonds in both acyl chains). The distribution is dominated by mono- and di-unsaturated diacylglycerols but polyunsaturated lipids (6-15 unsaturations) increase markedly in deep aquifer fluids.

Stable carbon isotopic compositions of lipids point to an autotrophic origin of microbial biomass. Due to the high *in situ* concentration of CO_2_ (at saturation level throughout the geyser [10]), maximum fractionation by carbon-fixing microorganisms in the geyser can be assumed [38]. Based on this, and the known fractionation (ε) of carbon fixed via different pathways [39], it is plausible that the majority of archaeal lipids were probably synthesized via the Wood Ljungdahl (WL, reductive acetyl-CoA, εDIC-lipid > 30‰) pathway from DIC, with εDIC-lipid of 38.3-53.9‰ observed in phytane derived from archaeol-based IPLs (Fig. 2B). This is in accordance with previous investigations of Crystal Geyser, which reported dominance of Altiarchaeoata in the deepest aquifer [10]. Altiarchaeoata fix carbon via a variant of the WL pathway with a fractionation of ~63‰ relative to DIC [40] (assuming εDIC-CO_2_ as ~10‰ at 15 °C calculated after ref. [41]). The observed εDIC-lipid values for archaeal lipids in many samples are below the maximum theoretical fractionation, implying that archaea in Crystal Geyser are not exclusively autotrophic but also take up isotopically heavier organic carbon. One likely source is archaeal utilization of organic carbon fixed by bacteria via the Calvin-Benson-Bassham (CBB) and reductive tricarboxylic acid cycle (rTCA) cycles, which would be more enriched in ^13^C than carbon fixed via the WL pathway. The degree of heterotrophic uptake by archaea can be approximated using a mass balance calculation involving mixtures of carbon with (i) the maximum theoretical fractionation for autotrophic archaeal carbon fixation via the WL pathway and (ii) the observed δ13C values of bacterial lipids. This calculation would imply that archaea are predominantly autotrophic in deep groundwater (up to 65% of the biomass carbon fixed through WL pathway), but in the intermediate and shallow groundwater form up to 80% of their biomass by taking up bacterial organic carbon fixed through the CBB and rTCA cycles (Fig. 2b).

Bacterial lipids display the carbon isotopic fractionation expected from the CBB cycle relative to CO_2_ (εCO_2_-lipid of 20.9-28.8‰ observed vs. 30‰ theoretical) and not that expected from fixation via the rTCA cycle (εCO_2_-lipid < 12‰ theoretical). Sequences encoding the CBB pathway are fairly abundant in the ecosystem throughout the recovery phase [10] and likely contributed to the bacterial lipid pool of samples collected during that period. This agrees with previous genomic findings that identified several highly active iron-oxidizing *Gallionella* species carrying this pathway [10]. Importance of *Gallionella* for *in situ* carbon fixation is further indicated by the association of organic carbon with fossilized *Gallionella* cells in post-depositional iron concretions of the Navajo sandstone [36]. However, genomic analyses suggested that one of the most abundant organisms in the shallow aquifer (*Sulfurimonas* sp.) fixes carbon via the rTCA cycle [10]. From mass balance calculations using the observed and theoretical fractionations, we estimate that carbon fixed via the rTCA cycle contributes as little as 6% to the bacterial biomass in the deep and intermediate aquifer but up to 51% of the biomass in the shallow aquifer (Fig. 2b). Overall, the observed carbon isotopic composition of the bacterial lipids could be explained as the result of a mixture of *Sulfurimonas*-derived lipids and lipids formed via the CBB pathway.

### Degree of unsaturation of bacterial IPL changes with groundwater source depth

Using in-depth analyses of IPLs we tracked the abundance of IPL-bound bacterial unsaturated FAs across the eruption cycle. The unsaturations presumably correspond to double bonds but due to the mode of detection, we cannot strictly rule out cycloalkyl groups found in FAs of some bacteria [47], although typically not in higher numbers than one per fatty acid. Interestingly, the relative abundance of highly unsaturated FAs correlated with the groundwater depth source (Fig. 2c). The cumulative abundances of IPLs with one or two double bond equivalents in their fatty acid side chains were fairly consistent throughout the cycle, indicating little variation between the different groundwater sources. However, IPLs with seven or more unsaturations, i.e., at this high number presumably double bonds, were relatively abundant during the first phase, where groundwater from intermediate depths is sourced. These lipids were even more abundant during the middle phase, during which groundwater derives from the greatest depth, and almost undetectable in samples collected in the final shallow groundwater eruption phase. One explanation for elevated abundance of polyunsaturated lipids is their derivation from eukaryotes [48]. The tentatively identified of DGCC-type (1,2-Diacylglyceryl-3-O-carboxyhydroxymethylcholine) betaine lipids is unprecedented in bacteria and supports the presence of Eukaryotes in the ecosystem, although the pathway for generating these lipids remains unknown [49]. In general, Eukaryotes have been found in the geyser [50], primarily in a sample of decayed wood added to the geyser conduit, and they have been detected by rpS3 analysis in the current study. However, they are not very abundant, and fluctuate heavily throughout the cycle (Fig. S6). The most likely explanation for the presence of highly unsaturated FAs is their origin from bacteria adapted to high pressures in the deeper subsurface. Incorporation of double bonds in bacterial FAs is a well-known mechanism that increases membrane fluidity at high pressure and low temperature [51, 52]. As temperature remained nearly constant at around 17 °C (Fig. 2), high fatty acid unsaturation could represent an adaptation to the high pressures faced by indigenous bacterial communities in the intermediate and deep aquifers, supporting a direct link between groundwater sources and lipid profiles.

### Predicting linkage of IPLs to uncultivated organisms

We detected 295 different IPLs in the 14 lipidomes but a strict organism-lipid relation in this relatively complex community was unresolved. The relative abundance of archaeal 16S rRNA genes was previously shown to correlate well with the relative abundance of ether lipids [53]. Moreover, various lines of evidence suggest that recycling of lipids may be a common strategy utilized by energy-starved archaea in the subsurface [54–56]. In the current study, we used a time series of 14 metagenomic and coupled lipidomic data sets to establish correlations between marker gene abundances and IPLs. Based on this analysis, we tested for evidence for the assignment of lipids to organisms. Specifically, relative abundance patterns of individual organisms were correlated with the relative abundance of the 295 IPLs (only organisms and lipids were considered if they were identified in at least seven out of fourteen samples). Lipids were also co-correlated with other lipids and primary and secondary lipid assignments were investigated via a network analysis (Fig. 3). Table 1 summarizes the detected correlations between lipids and organisms: 44 primary lipids correlate significantly with 22 different marker genes (organisms) and 63 secondary lipids.

**Fig. 3:**
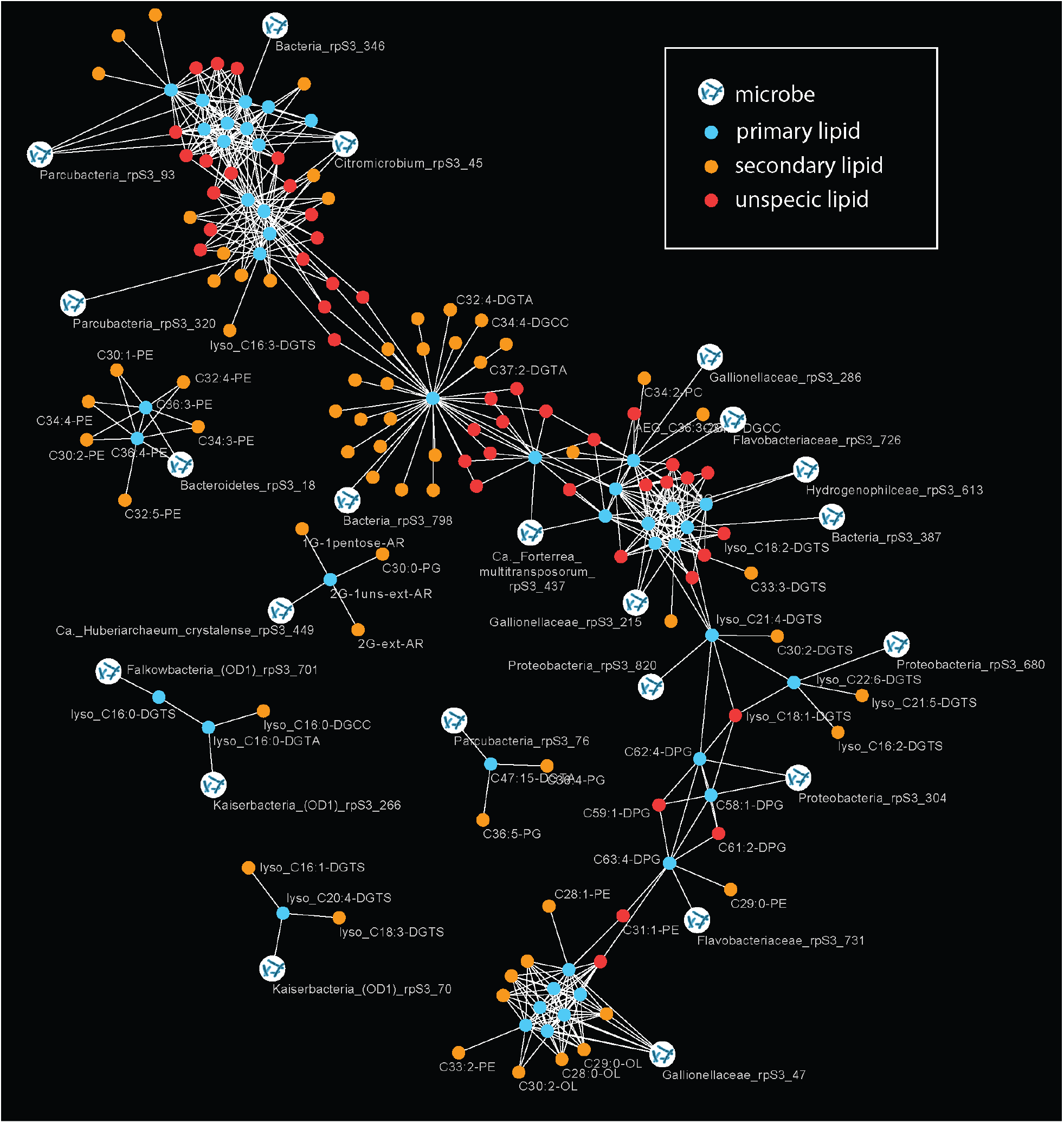
Correlation network analysis of relative abundances of organisms (rpS3 genes) and relative abundance of IPL signatures. The primary lipids were defined based on a direct correlation of their relative abundance with rpS3 gene abundance (Bonferroni-corrected p-value < 0.005). Secondary lipids showed a significant correlation with primary lipids and are indicative of a biological connection between the lipids (e.g. lipids from microbial symbionts or co-correlated organisms). Unspecific lipids shared primary lipids with different organism assignment. Due to visual limitations only few IPL names are displayed in the figure; all organisms to lipid correlations are provided in Table 1.

**Table 1:**
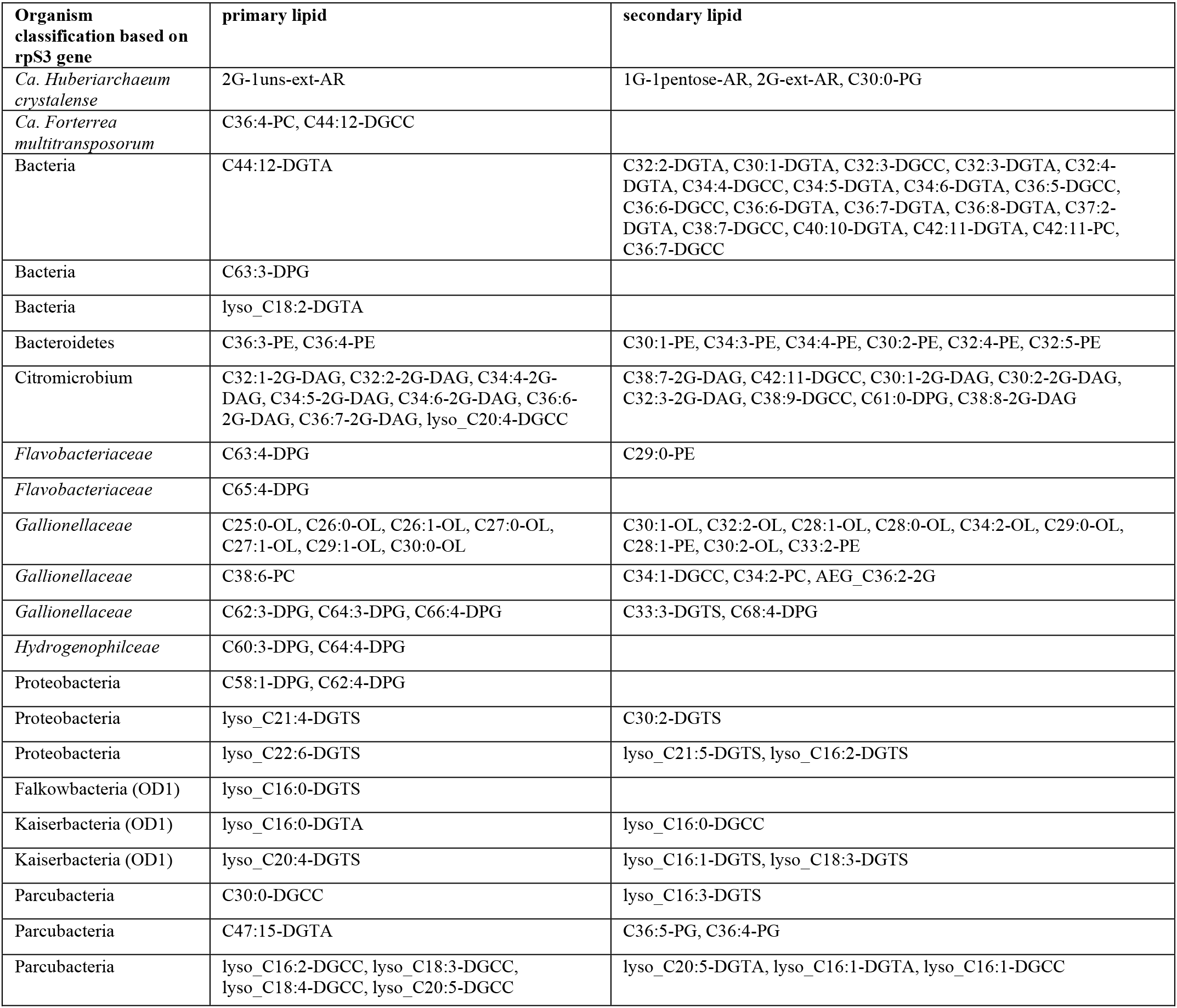
Correlation of rpS3 gene abundances from metagenomic read mapping with relative abundance of IPL signatures across samples. Primary lipids are direct correlations, secondary lipids are those that correlated with primary lipids.

It is important to note that all significantly correlating ether-based isoprenoid lipids were assigned to archaea (*Ca*. Huberiarchaeum crystalense) as this provides confidence in the correlation-based approach. However, it is unclear whether correlation of the main lipid of the *Ca*. H. crystalense and one bacterial lipid is a spurious covariation or if this represents assimilation of a bacterial membrane lipid by archaea (*Huberiarchaeum* did not correlate with that bacterial lipid; Table 1). Of particular interest were the lipids of Altiarchaeota, since these had been characterized earlier [40]. These previously detected lipids, including hexose-pentose archaeol (1G-1pentose-AR; for mass spectrometric identification see Fig. S4) and dihexose extended archaeol (2G-AR), were the most abundant archaeal lipids in the current study but most abundances showed little correlation with the Altiarchaeota abundances. On the one hand, this might be due to the presence of multiple different strains of *Altiarchaeum sp*. in the samples (based on rpS3 genes; Fig. S7), which can harbor different lipid profiles as shown previously [57]. On the other hand, the main archaeal IPL (2G-AR) was also present on the post 0.2-µm filter but *Altiarchaeum sp*. DNA was not (based on rpS3 genes). This indicates the lysis of *Altiarchaeum sp*. during the filtration process, possibly due to oxygen stress, a resistance that *Altiarchaeota* in Crystal Geyser apparently do not possess [40]. Nevertheless, some IPL signatures (e.g., 2G-ext-AR) showed a significant correlation with the sum of rpS3 abundances of all *Altiarchaeum sp*. in the sample, supporting the abovementioned assumptions (Fig. S7).

We detected one low abundance archaeal lipid, an unsaturated variant of 2G-ext-AR (2G-1uns-ext-AR), which had not been identified in Altiarchaeota. This may be a previously unrecognized membrane component of Altiarchaeota or derived from another archaeon. Its abundance correlated only weakly with other Altiarchaeota lipids but highly significantly with the abundance of Huberiarchaeum, thus it may derive from this organism. Huberarchaeota are the second most abundant archaea after Altiarchaeota in this ecosystem and they are predicted to have the genes required to synthesize lipids from scavenged isoprenoids [10]. The molecular structure of 2G-1uns-ext-AR differs by only one double bond from the Altiarchaeota lipid 2G-ext-AR, so Huberarchaeota may largely derive its lipids from Altiarchaeota, which was suggested to be its host [10]. The relative abundance of 2G-1uns-ext-AR correlated significantly with 2G-ext-AR, highlighting the potential biological meaning that can be inferred from IPLs, whose abundances do not correlate with certain organisms but with certain lipids instead. Given the confident assignment of 2G-1uns-ext-AR to Huberarchaeota, we used the p-value for that assignment as a conservative correlation p-value for further predictions (Bonferroni-corrected p-value <0.005), which are presented in Table 1.

### Lysolipids and Candidate Phyla Radiation bacteria

In order to investigate lipids of bacteria from the Candidate Phyla Radiation (CPR [33]), we analyzed the IPLs of a small cell size-fraction collected on a 0.1-µm pore-size filter after 0.2-µm pre-filtration. Based on the corresponding metagenome, the sample contained 186 different organisms, 165 of which were classified as CPR based on rpS3 sequences and one low abundant organism was classified as a member of the DPANN radiation (*Cand*. Huberiarchaeum crystalense). Surprisingly, the most abundant organism in the sample based on metagenomics was a *Sulfurimonas*, which apparently passed through the 0.2 µm filter (read mapping-based coverage in 0.2 µm filter was 8.4 in the corresponding 0.1 µm filter 1081.9). We identified 72 different IPLs in the post-0.2-µm sample, none of which were bacterial ether lipids. Consequently, the CPR organisms in this sample must possess fatty acid-based lipids. This is important because the composition of lipids of CPR bacteria is unknown. Interestingly, 22 of the 72 lipids (31%) were lysolipids, all of which contained betaine headgroups (for structural characterization see Fig. S3). By contrast, these lipids constituted only 18% across the entire sample set. Usually bacteria only contain a small fraction of lysolipids, e.g., *Sulfurimonas* has been reported to only contain a single lysolipid with ~4% abundance [59]. Further, the abundances of several CPR bacteria also correlated significantly with the abundance of specific lysolipids (Table 1).

We further investigated cells that passed through a 0.2-µm filter and were collected onto a 0.1-µm filter for lysolipids. Previous metagenomic sequencing analysis of the selected sample (CG10_big_fil_rev_8_21_14_0.10; [10]) showed a high abundance of CPR (rank abundance curve in Fig. S8). To further confirm these observations, we performed FTIR analysis of the cells (Fig. 4a) and compared the results against a set of reference spectra (Fig. S9). For the first PCA in the 3050-2800 cm^-^1 spectral region dominated by the aliphatic chains of the lipids, ~85% of the spectral variance is explained by the first five loading vectors (Fig. 4b). Here, the first loading vector contains the 55% of the variance, with features that are similar to lyso-phosphatidylcholine; with the asymmetric stretching of the CH_2_ peak centered at 2918 cm^-^1 whereas the phosphatidylcholine is centered at 2922 cm^-^1 (Fig. 4). The corresponding heatmap of the PC1 scores (Fig. 4c) shows the presence of hotspots, a few microns in diameter. The remaining 2, 3 and 4 loading vectors, which explain 18%, 10% and 2% of the variance respectively, show different CH_3_ to CH_2_ ratios. We noticed that the peak position of the asymmetric CH_2_ vibration for all the four loading vectors (1 to 4) is red-shifted towards a lysolipid-like absorption feature. In contrast, although the fifth loading vector accounts for only 1% of the variance, its spectral features can be assigned to highly branched and unsaturated lipids similar to those of archaea (Fig. 4b-c; refs. [12, 59]; see supplementary material for additional results). This agrees with the presence of DPANN archaea as the second most prominent group of organisms in this sample based on metagenomic profiling (Fig. S8). The combination of the detailed analysis of the IPLs and infrared imaging of two independently sampled small cell fractions led us to conclude that a substantial fraction of some CPR cell membranes consists of lysolipids.

**Fig. 4:**
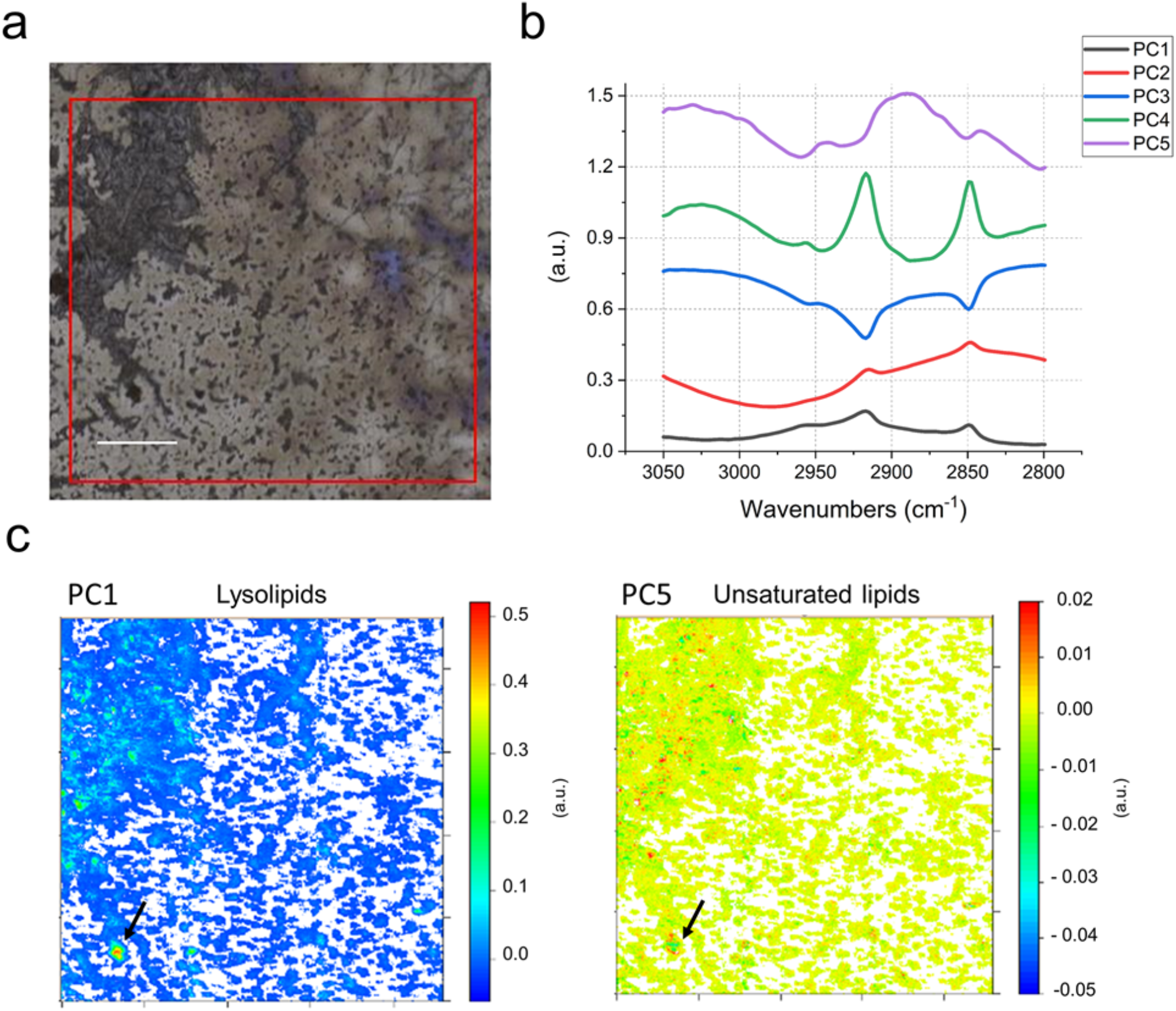
FTIR analysis of a small cell size fraction (post 0.2-µm filter collected onto a 0.1-µm filter). a) Field of view in FTIR, 1×1 mm. b) First five PCA loadings describing ~90% of the variance. c) False color maps representing PCA scores PC1 and PC5. By comparison of the spectral features of the loadings and the reference spectra in Fig. S1 allow assignment of PC1 to lysolipids and PC5 to unsaturated lipids. The arrows point to a hotspot of cells indicating a particularly high distribution of lysolipids (PC1), surrounded by several smaller hotspots of unsaturated lipids (PC5). Given the micrometric lateral resolution of the image (each pixel is 2.6 µm) it is possible to hypothesize that there is a small group of cells in the hotspot area, which is characterized by distinct membrane lipid composition. This can also be observed in other spots throughout the measured biomass. Scale bar 200 µm.

Genome-resolved metagenomics generated 206 new genomes from the entire sample set. Together with 1,215 previous genomes [10, 30], our dataset included 675 genomes of CPR bacteria that were used to comprehensively investigate their potential for lipid biosynthesis (accessible through https://ggkbase.berkeley.edu/genome_summaries/1491-Bacterial_membrane_lipids_AJP). We found that the CPR genomes do not encode for any known, complete bacterial lipid biosynthesis pathway, yet CPR bacteria are known to have a cytoplasmic membrane based on cryogenic-transmission electron microscopy studies [12]. Interestingly, some members of the Nealsonbacteria phylum (Parcubacteria superphylum) have near-complete pathways for FAs and phospholipid synthesis. They possess some homologs of the FA synthase type II (FAS-II), the main FA biosynthesis pathway in most bacteria. However, they lack the FAS-related acyl carrier protein (ACP) processing machinery (acyl carrier protein synthase and malonyl-CoA:ACP transacylase). ACP is a peptide cofactor that functions as a shuttle that covalently binds all FA intermediates. Although they lack key genes for FA synthesis, we cannot rule out this group could potentially synthetize FAs by an ACP-independent pathway, as suggested for some archaea ([60]). We also searched theses genomes for genes coding for glycerol-3-phosphate (G3P) dehydrogenase, an enzyme responsible for the stereochemistry of the glycerol units of their membrane lipids, and acyl-ACP transferases responsible for the formation of ester bonds between FAs and G3P backbone in phospholipid synthesis. There are two families of acyltransferases responsible for the acylation of the C1-position of the G3P. The PlsB acyltransferase primarily uses ACP end products of FA biosynthesis (acyl-ACP) as acyl donors. The second family involves the PlsY acyltransferase and is more widely distributed in Bacteria. PlsY uses as donor acyl-phosphate produced from acyl-ACP by PlsX (an acyl-ACP:PO4 transacylase enzyme). The acylation in the C2-position of the G3P is carried out by the 1-acylglycerol-3-phosphate O-acyltransferase (PlsC). Screening the Nealsonbacteria genomes, we did not detect any homologs of the first family of acyltransferase, PlsB. However, we identified PlsY and PlsC, but not PlsX. Absence of PlsX raises the question of the enzyme or mechanism for production of acyl-phosphate needed to activate PlsY. Overall, mechanisms or enzymes that produce and/or require ACP were not identified in CPR genomes in this study. Even though this finding opens the possibility for the presence of ACP-independent pathways for FA and/or lipid synthesis in these CPR bacteria, we cannot conclude with confidence that few of these organisms can synthesize lipids *de novo*. Thus, we suggest that most CPR bacteria derive their membrane lipids, including lysolipids, from co-existing bacteria. Given the small cell size of CPR, lysolipids may be preferred due to their role in reducing membrane curvature stress (e.g., [61]). As lysolipids can form during lipid breakdown (e.g., mediated by phospholipase A [62]) and can be taken up by other bacteria [63], their utilization by CPR may indicate uptake from degraded bacterial biomass or direct derivation from host cells.

### Model of lipid transfer in the community and conclusions

Our approach combined detailed metagenomics with whole community lipidomics and infrared spectroscopy and was informed by isotopic measurements that were constrained by detailed understanding of the geological context. The objective was to probe the carbon cycle within the subsurface microbial ecosystem, particularly the source of fixed organic carbon, but also to investigate evidence for its redistribution into other organisms, especially putative symbionts. Although sample limitation resulted in a lower resolution of isotopic analyses compared to metagenomics, carbon isotope systematics of archaeal and bacterial lipids confidently support the metagenomic predictions that microbial biomass is mostly of autotropic origin in all aquifers sampled. Particularly, our results provide evidence that predicted autotrophs were fixing CO_2_ *in situ*, using the WL (Altiarchaeum), rTCA (*Sulfurimonas*), and CBB cycles (*Gallionella*).

Using lipidomics and infrared spectroscopy on size-fractionated cells, we demonstrate that CPR bacteria with small cell size possess FA-based IPLs, although the corresponding genomes do not encode for a known pathway to synthesize them. Similarly, Huberarchaeota, potential symbionts of Altiarchaeota, were predicted to possess altered archaeal lipids related to those of their putative hosts. Our results support the notion that organisms of the CPR and DPANN radiation do not only scavenge (or symbiotically receive) molecular building blocks or even intact lipids from other bacteria and archaea but also use the corresponding lipids and introduce modifications (Fig. 5).

**Fig. 5:**
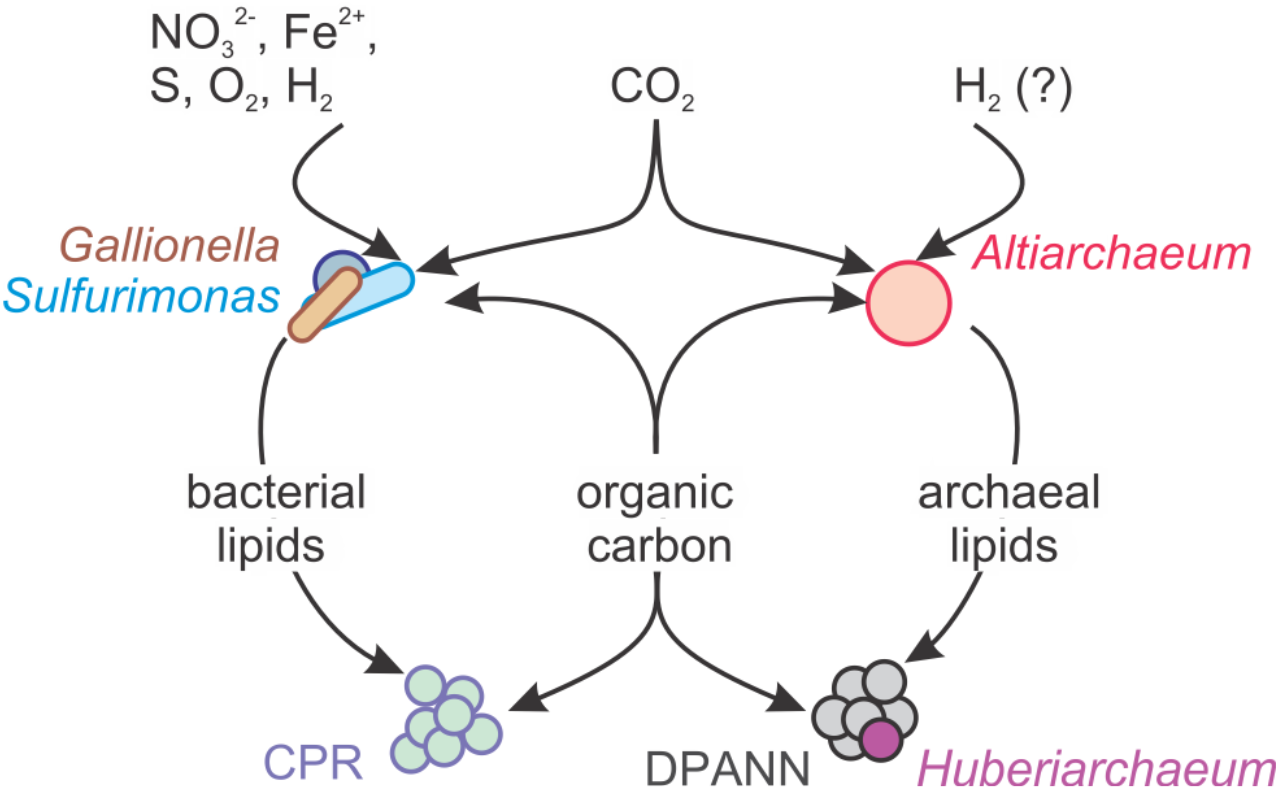
Model for the acquisition and redistribution of carbon and lipids in the deep subsurface ecosystems of the Colorado Plateau (USA) accessible through Crystal Geyser.

## ACKNOWLEDGEMENTS

We are grateful to Susan Spaulding, Christopher T Brown, and Karthik Anantharaman for assistance with sampling. We thank Julius Lipp for providing archaeal lipid standards and supporting lipid analyses. This study was funded by the Sloan Foundation (“Deep Life,” grant no. G-2016-20166041). AJP was also supported by the Deutsche Forschungsgemeinschaft (DFG PR 1603/1-1) and acknowledges funding by the Land Nordrhein-Westfalen (Nachwuchsgruppe Dr. Alexander Probst). Lipid extractions and analyses at the University of Bremen were supported by the Deutsche Forschungsgemeinschaft through the Gottfried Wilhelm Leibniz Program (award to KUH; Hi 616-14-1) and a project grant to JFB, AJP, and KUH by the “Deep Life Community” of the “Deep Carbon Observatory”, which is supported by the Alfred P. Sloan Foundation. The work conducted by the U.S. Department of Energy Joint Genome Institute, a DOE Office of Science User Facility, and by DOE’s Berkeley Synchrotron Infrared Structural BioImaging Resource at the Advanced Light Source are supported under Contract No. DE-AC02-05CH11231.

